# Efficient knock-in method enabling lineage tracing in zebrafish

**DOI:** 10.1101/2022.07.15.500272

**Authors:** Jiarui Mi, Olov Andersson

## Abstract

The CRISPR-Cas9 system aids generation of knock-in zebrafish lines, but it has been hard to integrate large constructs and avoid disrupting the targeted genes. Here we devised a 3’ knock-in strategy of PCR-amplified dsDNA, which coded for fluorescence proteins and Cre recombinase in frame with the endogenous gene but separated from each other by self-cleavable peptides. Primers with 5’ AmC6 end-protections generated improved PCR amplicons harboring either short or long homologous arms, which were co-injected with pre-assembled Cas9/gRNA ribonucleoprotein complexes for early integration. We targeted four genetic loci (*krt92*, *nkx6.1, krt4,* and *id2a*) and generated ten knock-in lines, which function as reporters for the endogenous gene expression. The knocked-in iCre or CreERT2 were used for lineage tracing, which suggested *nkx6.1^+^* cells are multipotent pancreatic progenitors that gradually restrict to bipotent duct; while *id2a^+^* cells are multipotent in both liver and pancreas and gradually restrict to ductal cells. Additionally, hepatic *id2a^+^* duct show progenitor properties upon extreme hepatocyte loss. Thus, we present an efficient knock-in technique with widespread use for both cellular labelling and lineage tracing.

## Introduction

With the advent of genome editing tools, the CRISPR-Cas9 system has become the most popular method in the generation of knock-in animal models (1, 2). Multiple knock-in strategies have been introduced and optimized for the construction of non-human primate (3, 4), mouse (4, 5), and zebrafish models (6, 7). In zebrafish, the knock-in methods vary in terms of the targeting regions (such as 5’ non-coding region, exon, intron, or 3’ end) (8–10), DNA double-stranded break repair mechanisms (homology dependent pair [HDR] or non-homologous end joining [NHEJ]) (11, 12), the type of donor templates (13, 14), the injection of Cas9 protein or Cas9 mRNA, and the use of drugs in promoting HDR (15, 16).

In zebrafish, the 5’ knock-in methods have been intensively investigated in the locus upstream of the ATG using donor plasmids containing *in vivo* linearization site(s) (11,12,17–20). The single linearization site upstream of the insertion sequence can facilitate the NHEJ-mediated integration (25411213, 28713249, 25293390); while researchers also tried to introduce long homologous arms (HAs) flanked by two *I-*SceI/gRNA recognition sites to induce HDR (27003937). Although fluorescence reporter lines, and even CreERT2 lines, have been generated by such methods (29435650), the wide applications of these methods are still hampered by the disruption of one allele of the endogenous gene as well as the cumbersome molecular cloning steps. The 3’ knock-in method has also been applied using circular plasmids as the donor, with either long or short homologous arms flanked by two *in vivo* linearization sites (14, 21–23). The advantage of 3’ knock-in is that it keeps the knock-in cassettes in-frame and maintain the functionality of the endogenous gene. However, in certain cases, a few amino acids in the C-terminus may be deleted when using the NHEJ strategy (30969163). Several studies reported that the HDR mediated 3’ knock-in efficiency is highly dependent on the length of the HAs (usually with higher efficiency using > 500 bp HAs), in particular when the gRNA targeting region is located upstream of the stop codon (27003937, 25411213). Nevertheless, one recent study showed that the introduction of short HAs in the donor plasmids flanked by two linearization sites can also enhance microhomology-mediated end joining (MMEJ) (29812974). However, such method is still limited in use due to the low scalability and the tedious construct preparation steps. Recently, intron-based and exon-based knock-in approaches have remarkably expanded the knock-in toolbox by targeting genetic loci beyond the 5’ or 3’ end (8-10,13,24-26). These methods mostly rely on the NHEJ method and the endogenous genes can be either destroyed or rescued (depending on whether or not the exon sequences downstream of insertion site are added into the donor). Given that all these methods are labour-intensive and limited in scalability, the development of a straightforward and efficient knock-in methodology is still warranted in the zebrafish field. Such method should not to be compromised by the common issues, including the disruption of one allele of the endogenous gene (knock-in/knock-out), the complex pipeline in molecular cloning, low efficiency of precise recombination, limited germline transmission rate, and high workload in screening for founders.

Recent studies in mouse and *in vitro* systems have demonstrated several approaches that improve HMEJ. The Tild-CRISPR (targeted integration with linearized dsDNA-CRISPR) strategy that used PCR-amplified or enzymatic-cut donors with 800 base pairs HAs has been successfully applied in generating knock-in mouse lines (29787711), indicating that nude double-stranded DNA (dsDNA) can serve as an effective donor in eukaryote embryos. Furthermore, 5’ modified dsDNA with short HAs (roughly 50 base pairs) demonstrated impressive knock-in efficiency in an *in vitro* culture system (31873222). In this study, the researchers systematically compared 13 modifications on dsDNA with gRNA targeting the 3′ untranslated region (UTR) of the GAPDH gene in HCT116 cells. The dsDNAs were synthesized by PCR amplifications with the modifications incorporated in the primers. It showed that C6 linker (AmC6) or C12 linker (AmC12) as well as moieties by adding on secondary modifications out-performed no C6/C12 linked modifications with a substantial increase of knock-in efficiency by more than five-fold. Although the mechanism is still undetermined, it is postulated that the 5’ modification can help prevent degradation and multimerization of the donor and circumvent stochastic NHEJ, indicated by less NHEJ events and random insertions.

Inspired by these previous efforts in improving knock-in efficiency (27), here, we introduce a straightforward CRISPR-Cas9 guided 3’ knock-in approach to generate zebrafish lines for cellular labelling and lineage tracing. We synthesized 5’ modified dsDNA with either short or long HAs as the donor by a simple PCR step with 5’ modified primers (AmC6). The donor templates code for two kinds of 2A peptides linking the endogenous gene product with a fluorescent protein and then with iCre/CreERT2. By co-injecting this type of donor with *in vitro* pre-assembled Cas9/gRNA ribonucleoprotein complexes (RNPs), we achieved very high percentage of mosaic F0 giving rise to germline transmission. Ultimately, we generated ten knock-in fishlines, demonstrating the high scalability. Our knock-in lines can precisely reflect the endogenous gene expression, as visualized by optional fluorescent proteins. Importantly, we also performed lineage tracing experiments using the knock-in iCre and CreERT2 lines to delineate cell differentiation paths in pancreas and liver development and injury models.

## Results

### A 3’ knock-in pipeline and the characterization of *TgKI(krt92-p2A-EGFP-t2A-CreERT2)*

With the aim to generate knock-in zebrafish lines for both cellular labelling and lineage tracing, we designed our vector templates encompassing a fluorescence protein and different Cre recombinase linked by two self-cleavable 2A peptide sequences (p2A and t2A) (Figure 1A). The insertion sequences were flanked by left and right long HAs (nearly 950 base pairs) on the basis of the GRCz11 reference genome archived in the Ensembl database. Next, we used pairs of 5’ AmC6 modified primers to amplify dsDNA with the insertion sequence flanked by either long or short HAs by PCR. Subsequently, we co-injected the PCR product, which serves as the direct donor, together with *in vitro* pre-assembled Cas9/gRNA ribonucleoproteins (RNPs) into the one-cell stage zebrafish embryos (of the TL strain). In the next few days, we sorted out the F0 with high percentage of mosaicism based on the fluorescence in the expected cell types and raised them up. The adult F0 were then outcrossed with wild type fish to screen for founders. To test the validity and efficiency of this pipeline, we first selected an epithelial marker, *krt92*, as it facilitated the identification of mosaic F0 by a simple detection of fluorescence in the skin.

**Figure 1.**
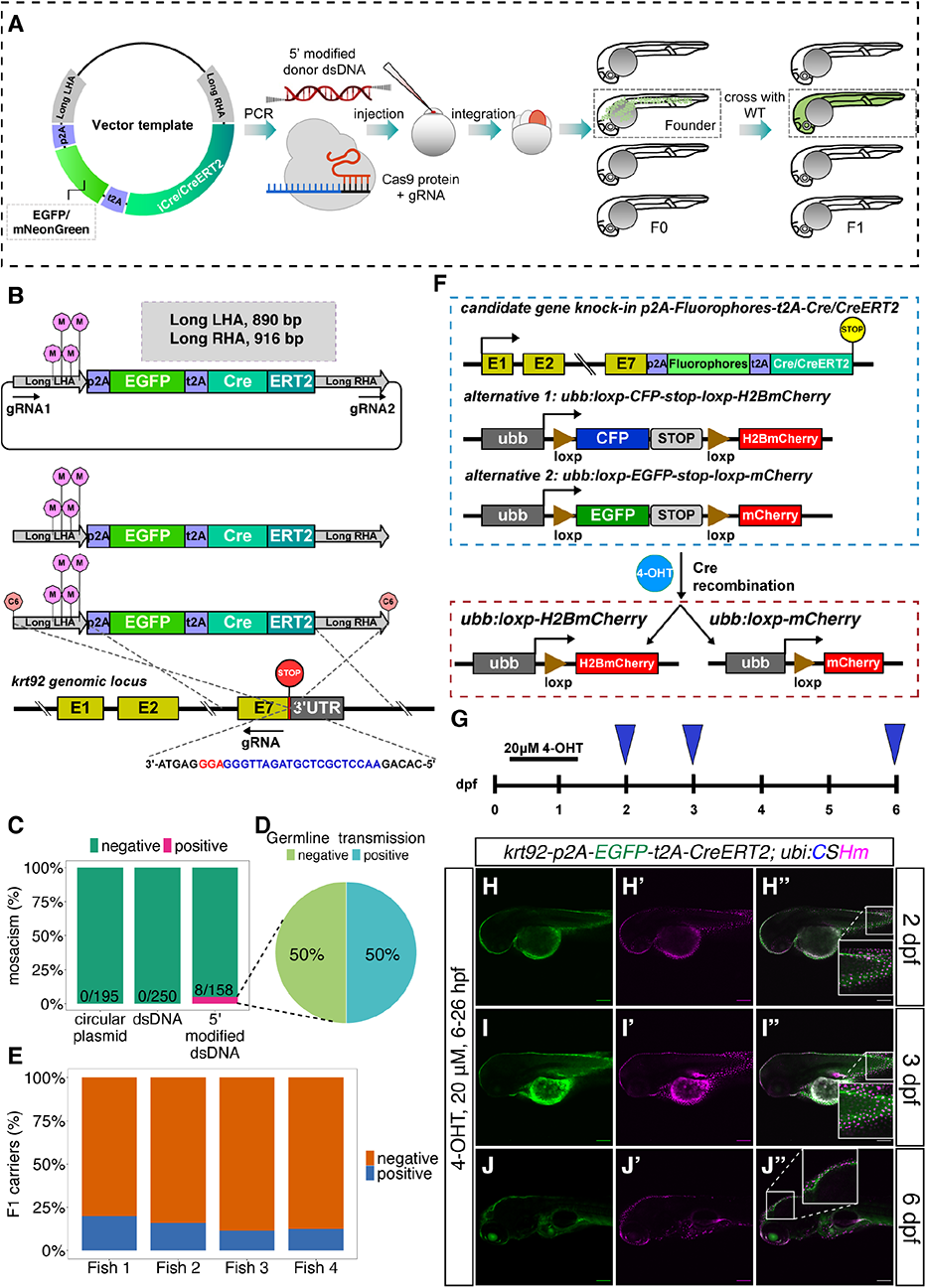
Knock-in pipeline and characterization of knock-in line at the *krt92* locus. Schematic representation of the 3’ knock-in pipeline with 5’ modified dsDNA as the donor. The iCre indicates improved Cre. (B) The design of the template vector, PCR-amplified dsDNA with 5’ modifications, and the gRNA sequence for the construction of *TgKI(krt92-p2A-EGFP-t2A-CreERT2)*. The gRNA1 and gRNA2 indicate *in vivo* linearization sites. The purple lollipops in the middle of long LHA indicate synonymous point mutations on the left HAs. The pink lollipops at the end of dsDNAs indicate 5’ AmC6 modification. The nucleotide sequence in blue indicates gRNA; while the nucleotide sequence in red indicates the PAM sequence. (C-E) Summary statistics of *krt92* knock-in efficiency using different donors, including the percentage of injected F0 with at least 30% fluorescence labelling in the skin (C), the percentage of adult F0 giving rise to germline transmission (D) and the percentage of F1 siblings from four different founders (Fish 1-4) carrying knock-in transgenes (E). (F) The scheme of lineage tracing strategy for the iCre or tamoxifen-inducible Cre knock-in lines. There are two alternatives for the color switch using the Cre responder lines, the option 1 contains *ubb:loxp-CFP-stop-loxp-H2BmCherry* transgene (abbreviated as *ubi:CSHm)*, while option 2 contains *ubb:loxp-EGFP-stop-loxp-mCherry* (abbreviated as *ubi:Switch)*. Cells with Cre recombination will ubiquitously express H2BmCherry or mCherry. (G-J) Temporal labelling with 20 μM 4-OHT treatment at 6-26 hpf in *TgKI(krt92-p2A-EGFP-t2A-CreERT2);Tg(ubi:CSHm)* line (G) and representative confocal images at 2 dpf (H-H’’), 3 dpf (I-I’’), 6 dpf (J-J’’). Skin cells with Cre recombination after 4-OHT treatment were labelled with H2BmCherry. The insets are magnified views showing the expression pattern of two fluorescent proteins. Scale bars = 200 μm.

For the *krt92* locus, the dsDNA donor and gRNA sequence are shown in Figure 1B and Figure EV1. We selected one gRNA 20 base pairs upstream of the stop codon. To circumvent the cleavage of the donor, and keep the endogenous amino acid sequence intact, we incorporated several synonymous point mutations on the left HA (Figure EV1). We used long HAs on both sides to enhance the annealing of the sequences. With 5’ modified dsDNA injection, we observed that 8 out of 158 (5.1%) injected embryos showed fluorescence in approximately a third of the skin, suggesting early integration (Figure 1C). Among them, four fish were identified as founders by screening more than 200 F1 embryos from each fish (Figure 1D). The proportion of the F1 generation that carried the transgenes ranged from 11.5% to 20% (Figure 1E).

To validate the functionality of Cre recombinase, we crossed the *TgKI(krt92-p2A-EGFP-t2A-CreERT2)* with the responder line *Tg(ubb:loxP-CFP-STOP-Terminator-loxP-hmgb1-mCherry)* (abbreviated as*Tg(ubi:CSHm)*) (Figure 1F). The cells with *krt92* expression during the 4-hydroxytamoxifen (4-OHT) treatment were expected to have Cre recombinase translocated to the nucleus to conduct recombination. Therefore, all the progenies of *krt92^+^* cells after the recombination would express H2BmCherry as it is directly driven by *ubiquitin B* promoter (30). The H2BmCherry signal was detected in skin cells after 4-OHT administrated at 6 hours postfertilization (hpf) (Figure 1G-J, Figure EV2). We also observed that various proportions of the intestinal cells were fluorescently labelled after 4-OHT treatments at different timepoints (Figure EV3). Although *in situ* hybridization for *krt92* is not feasible due to the high sequence similarity with other keratins, the fluorescence pattern matched the recent single-cell RNA-seq data of the zebrafish intestine, showing a widespread expression of across different intestinal epithelial cell types (36) (34301599). Lastly, we noticed that neither the circular plasmid with *in vivo* linearization sites nor the dsDNA without 5’ end protection can achieve successful integration (Figure 1B and C). In summary, the dsDNA with 5’ modification is an efficient donor for generating HDR-dependent knock-in zebrafish lines.

### The generation of *nkx6.1* knock-in lines using short or long homologous arms

Next, we aimed to knock-in donors at 3’ end of *nkx6.1*, which is a transcription factor essential in the development of the pancreas and motor neurons (39–40). We selected a gRNA spanning over the stop codon region, and used 5’ modified dsDNA with long HAs to generate *TgKI(nkx6.1-p2A-EGFP-t2A-CreERT2)* (Figure 2A). Two out of 1000 (0.2%) *nkx6.1-p2A-EGFP-t2A-CreERT2-*injected F0 embryos using long HAs showed detectable fluorescence signal in the spinal cord (Figure 3B).

**Figure 2.**
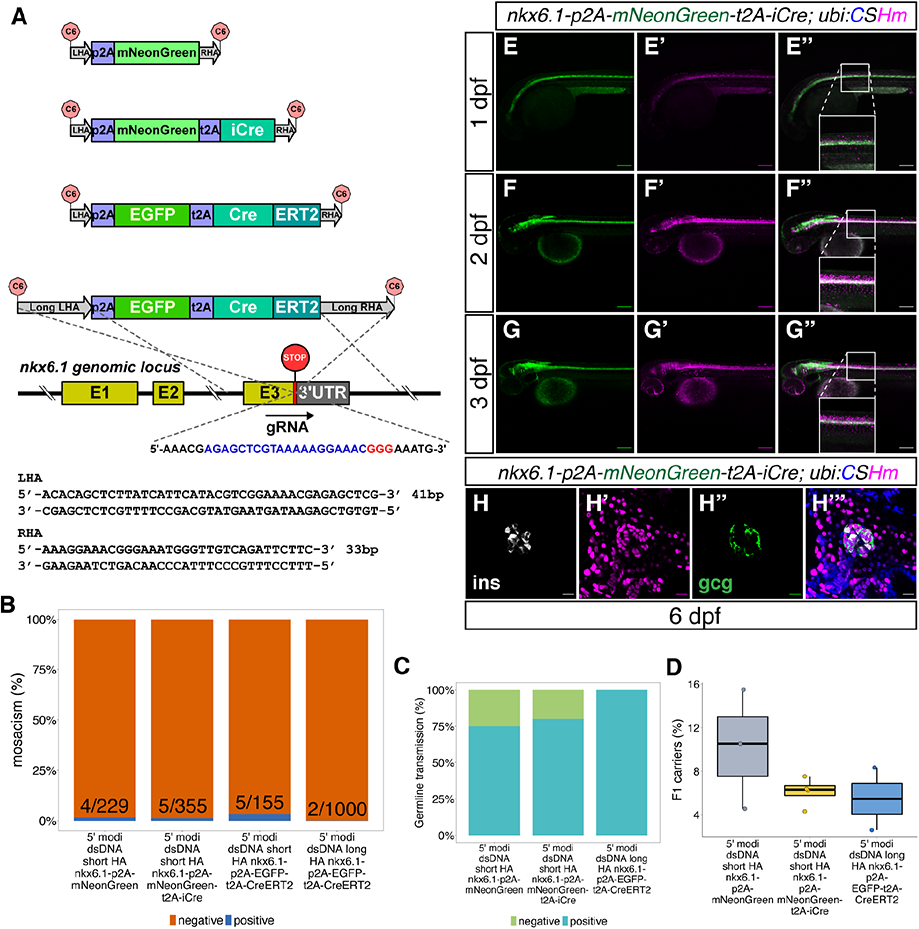
The design and characterization of knock-in lines at the *nkx6.1* locus. (A) The design of donor dsDNA template and gRNA sequence for the construction of knock-in lines at the 3’ end of the *nkx6.1* locus. The nucleotide sequence in blue indicates gRNA; while the nucleotide sequence in red indicates the PAM sequence. The bottom panel shows the sequence of LHA and RHA. (B-D) Summary statistics of *nkx6.1* knock-in efficiency, including the percentage of injected F0 with observable fluorescence labelling in the hindbrain and spinal cord (B), the percentage of adult F0 giving rise to germline transmission (C) and the percentage of F1 siblings carrying knock-in transgenes (D). (E-G) Representative confocal images of *TgKI(nkx6.1-p2A-mNeonGreen-t2A-iCre);Tg(ubi:CSHm)* at 1 (E), 2 (F), and 3 dpf (G). Cells expressing mNeonGreen indicate *nkx6.1^+^* cells; while all progenies of *nkx6.1^+^* cells were labelled with H2BmCherry. The insets are magnified views showing the expression pattern of two fluorescent proteins. (H) Representative confocal image of lineage tracing results in the principal islet of the pancreas in the *TgKI(nkx6.1-p2A-mNeonGreen-t2A-iCre);Tg(ubi:CSHm)* line at 6 dpf. Cells in white shown in H are β-cells with insulin staining; while cells in green in H’’ are α-cells with glucagon staining. Scale bars = 200 μm (E-G) or 20 μm (H).

**Figure 3.**
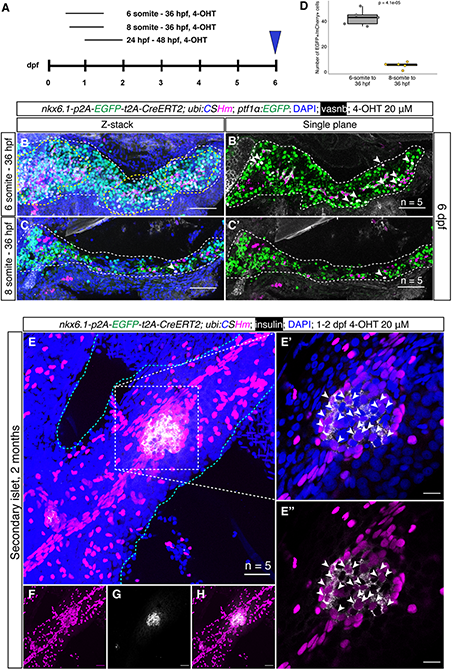
*nkx6.1*^+^ cells were gradually restricted to the duct and gave rise to secondary islets in the zebrafish pancreas. (A) Experimental timeline for temporal labelling using short-term and long-term lineage tracing. 4-OHT (20 μM) was added at the 6 or 8 somite stage and the treatment continued until 36 hpf for short-term lineage tracing. (B-B’ and C-C’) 4-OHT (20 μM) was added from 1-2 dpf and confocal imaging was performed at 60 dpf for the long-term lineage tracing (E-H). Representative confocal images of the pancreas at 6 dpf in *TgKI(nkx6.1-p2A-EGFP-t2A-CreERT2; ptf1α:EGFP);Tg(ubi:CSHm)*. The progenies of *nkx6.1*^+^ cells after 4-OHT treatment were H2BmCherry positive. The EGFP signals indicate acinar cells. Intrapancreatic ductal cells were demonstrated by membrane staining using the anti-vasnb antibody (shown in white). The region with yellow dashed line in B indicates H2BmCherry^+^/EGFP^+^ enriched regions. Arrowheads in B’, C, and C’ point to H2BmCherry^+^/EGFP^+^/vasnb^-^ cells, both indicating acinar cells from *nkx6.1^+^* cell origin. For each condition, we scanned 5 samples with 16-24 single-planes for the larval pancreata and 18-30 single-planes for the juvenile pancreata. The Z-stacked images were displayed (B and C) demonstrating a large number of mCherry^+^/EGFP^+^ cells with 4-OHT treatment starting from 6-somite stage; while the number of double positive cells are decreased with statistical significance. (D) The quantification and statistical results of EGFP/mCherry double positive cells with 4-OHT treatment starting at 6- and 8-somite stage. Two-tailed t-test was used for statistical analysis, with p-value < 0.05 considered as statistically significant. (E-H) Projection images of lineage-traced secondary islets in the *TgKI(nkx6.1-p2A-EGFP-t2A-CreERT2);Tg(ubi:CSHm)* zebrafish pancreas at 60 dpf. The progenies of *nkx6.1^+^* cells after the 4-OHT 1-2 dpf treatment were H2BmCherry positive. The selected area in the white dashed square in E was magnified in single plane (E’ and E’’). The cyan dashed lines outline the pancreas (E). Split channels of E are displayed for clarity (F-H). Arrowheads point to lineage-traced β-cells in the secondary islet co-stained with anti-insulin antibody. Scale bars = 80 μm (B-B’ and C-C’), 20 μm (E, F, G, H) or 10 μm (E’ and E’’), respectively.

As the donor plasmids with short HAs flanked by *in vivo* linearization sites were also applicable in zebrafish knock-ins by inducing MMEJ (14) (37–38), we converted to dsDNA with short HAs generated by a simple PCR step with primers containing 41 and 33 base pairs flanking the insertion sequence. We injected the dsDNA donor carrying *p2A-mNeonGreen* and *p2A-mNeonGreen-t2A-iCre* cassettes, and strikingly, we noted a dramatic increase of recombination efficiency, i.e. *nkx6.1-p2A-mNeonGreen-t2A-iCre* (in 5 out of 355 injected embryos) *or nkx6.1-p2A-mNeonGreen* (in 4 out of 229 injected embryos) (Figure 2B). The percentages of founders among these mosaic F0 were between 75% and 100% (Figure 2C); and 2.5% to 15.5% of the F1 siblings carried the knock-in transgenes (Figure 2D). For further comparison, we also injected *p2A-EGFP-t2A-CreERT2* dsDNA with short HAs and could identify 5 out of 155 (3.2%) injected embryos with detectable fluorescence. This indicated that short HAs outcompeted (more than 10-fold) long HAs at this locus (Figure 2B), i.e. when comparing different donors that all carried the 5’ AmC6 modification.

The iCre and CreERT2 function were characterized by the color switch in the offspring when crossed with *Tg(ubi:CSHm)*. We noticed that cells expressing *nkx6.1* (displayed by the green fluorescence) were located on the ventral side of the spinal cord; whereas H2BmCherry positive cells, which include all the progenies of *nkx6.1^+^* cells after the iCre recombination, resided in both the ventral and dorsal parts of spinal cord, suggesting a progenitor cell property of *nkx6.1^+^* cells in zebrafish spinal cord (Figure 2E-G, Figure EV4A-C). In addition, preceding immunostaining experiment using the *TgBAC(nkx6.1:EGFP)* reporter showed that *nkx6.1^+^* cells exist in both the dorsal and ventral buds of the pancreas at 17-48 hpf, indicating that they might be multipotent pancreatic progenitor cells (40–41); however, definitive evidence from lineage tracing experiments is still lacking. To further determine the *nkx6.1*^+^ cell lineage in the pancreas, we firstly performed the immunostaining for the fluorescent proteins in the knock-in lines and showed that *nkx6.1* specifically labelled the intrapancreatic duct in zebrafish larvae (Figure EV4D-F). Secondly, using the *nkx6.1* knock-in iCre line, we could trace back all three major cell types in the pancreas (acinar, ductal and endocrine cells) to *nkx6.1* lineage (Figure 2H-H’’’, Figure EV4G-I), suggesting that all these different cell types were indeed derived from the *nkx6.1^+^* cells.

### Short-term and long-term lineage tracing depicting the *nkx6.1* lineage

To further explore the cell fate determination in the zebrafish pancreas, we did lineage tracing experiment using *TgKI(nkx6.1-p2A-EGFP-t2A-CreERT2);Tg(ubi:CSHm)* with 4-OHT treatments at multiple timepoints (Figure 3A). The immunostaining at 6 dpf showed that both intrapancreatic ductal cells and a portion of acinar cells can be lineage traced when the 4-OHT treatment started at the 6-somite stage (Figure 4B and B’). In contrast, *nkx6.1* lineage-traced cells were mostly restricted to the intrapancreatic duct when the 4-OHT treatment started at the 8-somite stage (Figure 3C,C’ and D). These results pinpoint the exact timing of the early cell fate divergence between acinar cells and ductal/endocrine lineages.

**Figure 4.**
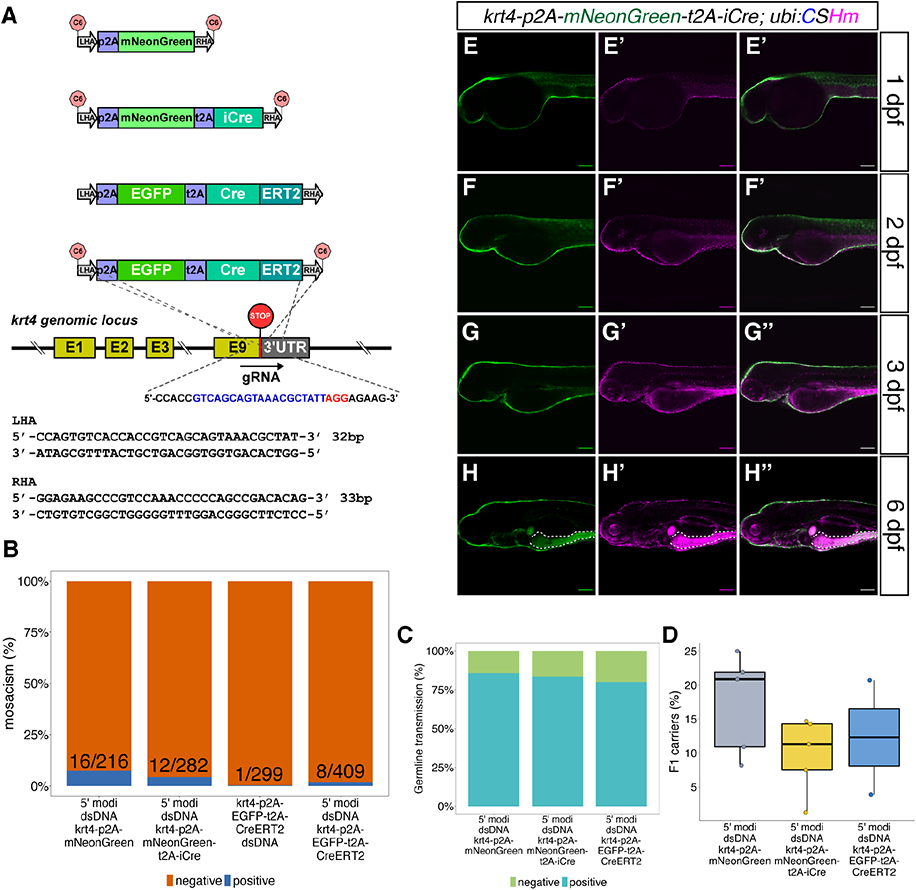
The design and characterization of knock-in lines at the *krt4* locus. (A) The design of donor dsDNA templates and gRNA sequence for the construction of knock-in lines at the 3’ end of the *krt4* locus. The nucleotide sequence in blue indicates gRNA; while the nucleotide sequence in red indicates the PAM sequence. The bottom panel shows the sequence of LHA and RHA. (B-D) Summary statistics of knock-in efficiency at the *krt4* locus, including the percentage of injected F0 with approximate one third of skin with fluorescence labelling. (B), the percentage of adult F0 giving rise to germline transmission (C) and the percentage of F1 siblings carrying knock-in transgenes (D). (E-H) Representative confocal images of *TgKI(krt4-p2A-mNeonGreen-t2A-iCre);Tg(ubiCSHm)* at 1 dpf (E-E’’), 2 dpf (F-F”), 3 dpf (G-G’’), and 6 dpf (H-H’’). Skin and intestinal epithelial cells (at 3 and 6 dpf) were broadly recombined and labelled with H2BmCherry. The white dashed lines outline the intestinal bulb (H-H’’). Scale bars = 80 μm.

Previous studies using transgenic lines based on *tp1*, which is a Notch-responsive element from the Epstein Barr virus mediating expression in the intrapancreatic duct (including *Tg(tp1:H2Bmcherry), Tg(tp1:venusPEST),* and *Tg(tp1:CreERT2)*), suggested that Notch-responsive intrapancreatic ductal cells can give rise to endocrine cells. These endocrine cells are mainly located in secondary islets and appear during growth or upon Notch inhibition (42–44). Additionally, previous immunostaining results in *TgBAC(nkx6.1:EGFP)* showed insulin/EGFP co-localization in adult zebrafish pancreatic tail region after β-cell ablation, suggesting latent duct-to-β-cell neogenesis is maintained in adulthood (40, 45). However, lineage tracing using *tp1:CreERT2* can only label at maximum 75% of the Notch-responsive ductal cells and traced very limited amount of endocrine cells, indicating that *tp1* was not an efficient tracer for ductal neogenesis (42, 46). To have a better understanding of duct-to-β-cell neogenesis, we performed a 2-month long-term lineage tracing experiment using *TgKI(nkx6.1-p2A-EGFP-t2A-CreERT2);Tg(ubi:CSHm)* with 4-OHT treatment at 1-2 dpf. We observed that nearly 65% of *ins^+^* cells in the secondary islets (residing along the large ducts) can be lineage traced (Figure 3E-H and Figure EV6D). Furthermore, we performed short-term lineage tracing experiments in the presence of a Notch inhibitor (LY411575) or a REST inhibitor (X5050), as previous studies described (29, 47), to demonstrate the neogenic potential of *nkx6.1^+^* intrapancreatic ductal cells in zebrafish larvae (Figure EV5). We found that more than 80% of *ins^+^* cells and around 75% of *gcg^+^* cells in the pancreatic tail can be lineage traced after three-day treatment (3-6 dpf) with LY411575 and X5050, respectively. These results, all together, suggested a latent progenitor property of *nkx6.1^+^* duct, one that can serve as the origin for endocrine cells.

Lastly, regarding the CNS, the temporal controlled experiments showed similar labeling pattern to the non-inducible Cre *nkx6.1* knock-in line, confirming that the *nkx6.1^+^* cells in spinal cord are also progenitors (Figure EV6A-C). Control experiment without 4-OHT treatment indicated no leakage problem with the *nkx6.1* knock-in CreERT2 line (Figure EV7).

### The generation of *krt4* knock-in lines using 5’ modified dsDNA with short homologous arms

Similar to *nkx6.1*, we identified one gRNA target spanning over the stop codon in *krt4* (Figure 2A). To assess the knock-in efficiency with a dsDNA donor using short HAs with 5’ modification, we amplified three different insertion sequences (*p2A-mNeonGreen*, *p2A-mNeonGreen-t2A-iCre*, and *p2A-EGFP-t2A-CreERT2*) with pairs of primers containing 32 and 33 base pairs homologous overhang at the 5’ ends by PCR (Figure 4A). Around 2.0% to 7.4% of injected F0 displayed at least one third of the skin with green fluorescence labelling at 1 dpf (Figure 4B). Additionally, more than 75% of these F0 were characterized as founders (Figure 4C). The F1 germline transmission rate across the three knocked-in lines ranged from 1.2% to 22.0% (Figure 4D). The iCre function was confirmed by crossing the *TgKI(krt4-p2A-mNeonGreen-t2A-iCre)* with *Tg(ubi:CSHm)*. We observed the H2BmCherry labelled cells in the skin and intestine (Figure 4E-H). However, *TgKI(krt4-p2A-EGFP-t2A-CreERT2);Tg(ubi:CSHm)* embryos showed mosaic leakage in the skin and intestine without 4-OHT treatment (Figure EV8A-C). Lastly, for systematic comparison, we also injected *p2A-EGFP-t2A-CreERT2* PCR products without end protection and observed that only 0.3% (1 out of 299) injected F0 achieved early integration based on the above criteria (Figure 4B), indicating that the 5’ modification can achieve around five-fold more efficient integration of dsDNA donors when using short HAs at this locus.

Lastly, as the *krt4* transgenics has been widely used for labeling skin epithelial cells (24415949, 24367659) (Figure EV8D), we performed a further characterization of *krt4* expression in the gut using HCR3.0 *in situ* hybridization and EGFP immunofluorescence in both *Tg(krt4-p2a-EGFP-t2a-CreERT2)* and *Tg(krt4:EGFP-Mmu.Rpl10a)* zebrafish larvae (Figure EV9). Notably, the *krt4 in situ* showed very strong signals in both the intestinal bulb and the hindgut. The green fluorescence in *krt4* knock-in line fully recapitulate the *krt4* endogenous expression (Figure EV9C and D); while the *krt4* transgenics displayed no signals in the gut (Figure EV9A and B), indicating that the *cis-*regulatory element cloned in the *krt4* transgenic line is unable to mimic the *krt4* expression and not sufficient to drive EGFP expression in the gut.

### The generation of *id2a* knock-in lines using short homologous arms

Next, we used similar strategies to knock-in *p2A-mNeonGreen*, *p2A-mNeonGreen-t2A-iCre*, and *p2A-EGFP-t2A-CreERT2* fragments into the 3’ end of the *id2a* locus using short HAs (Figure 5A). We found that 1.5% (2 out of 137), 0.8% (2 out of 252), and 3.0% (7 out of 240) injected embryos displayed strong fluorescence in the hindbrain, spinal cord and olfactory organs and were raised up to adulthood (Figure 5B). We observed high percentage of mosaic F0 with germline transmission (100%, 100%, and 71.4% respectively) (Figure 5C) and the percentage of the F1 generation carrying the transgene ranged from 3.2% to 29.0% (Figure 5D). We crossed the iCre and CreERT2 knock-in lines with *Tg(ubi:CSHm)* and observed similar pattern of mCherry signal as mNeonGreen in various tissues (hindbrain, dorsal side of spinal cord, pronephros, olfactory organ, and muscles), verifying the functionality of the Cre recombinases (Figure 5E-G, Figure EV10A-C). Control experiments suggested there is no leakage problem with the *id2a* knock-in CreERT2 line (Figure EV11).

**Figure 5.**
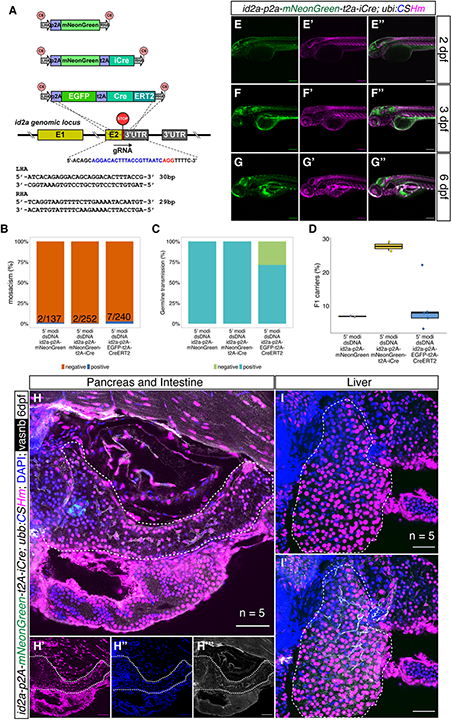
The design and characterization of knock-in lines at the *id2a* locus. (A) The design of donor dsDNA template and gRNA sequence for the construction of knock-in lines at 3’ end of the *id2a* locus. The nucleotide sequence in blue indicates gRNA; while the nucleotide sequence in red indicates the PAM sequence. The bottom panel shows the sequence of LHA and RHA. (B-D) Summary statistics of *id2a* knock-in efficiency, including the percentage of injected F0 with observable fluorescence labelling in the hindbrain, spinal cord and olfactory organs (B), the percentage of adult F0 giving rise to germline transmission (C) and the percentage of F1 siblings carrying knock-in transgenes (D). (E-G) Representative confocal images of *TgKI(id2a-p2A-mNeonGreen-t2A-iCre); Tg(ubi:CSHm)* at 2 dpf (E-E’’), 3 dpf (F-F’’), and 6 dpf (G-G’’). Cells that are mNeonGreen positive indicate *id2a-*expressing cells; whereas the progenies of the *id2a* lineage were mCherry labelled. (H-I) Representative confocal images of lineage tracing experiments in the zebrafish larval pancreata (H-H’’’), intestine (H-H’’’) and liver (I and I’) in the *TgKI(nkx6.1-p2A-mNeonGreen-t2A-iCre);Tg(ubi:CSHm)* line. We scanned 5 pancreata and livers, with 26 to 34 single planes imaged. Cells in cyan are α-cells based on anti-glucagon antibody staining. Scale bars = 200 μm (E-G) or 80 μm (H and I).

Previous studies have shown that *id2a* is important in the development of endodermal organs (e.g. liver and pancreas) (48–50) and retina (51). Therefore, we performed immunostaining of the knock-in *EGFP* line to visualize *id2a* expression in these tissues. In the zebrafish gut (intestinal bulb and hindgut), we observed a large number of *id2a^+^* cells showing overlapping fluorescence signals with *tp1:H2BmCherry*, suggesting that these cells are *id2a^+^* and have active Notch signaling (Figure EV10D, E). These double positive cells had apical membrane oriented towards the gut lumen, indicating that they were either sensory cells or secretory cells (Figure EV10F). This result was in line with two recent single-cell RNA-seq studies of the larval gut, proposing the subpopulation of Notch-responsive cells were *best4/otop2^+^* ionocytes (Figure EV12E-H) (36, 52), which regulate ion concentrations.

In the zebrafish pancreas and liver, the *id2a* knock-in reporter showed fluorescence in the intrapancreatic, a subset of extrahepatic and intrahepatic ductal cells, as well as in what we term the intermediate duct that branch out in between the extrahepatic and the intrahepatic duct (Figure EV10G-I). In the retina, the *id2a* reporter labelled a substantial number of retinal epithelial cells (Figure EV10K, L). Altogether, the *id2a* knock-in reporter closely recapitulated known expression domains of *id2a*.

### id2a tracing delineates developmental paths in the liver and pancreas

Previous fate mapping studies suggested that the pancreas and liver originate from common progenitors in the endodermal sheet, in which cells close to the midline are prone to differentiate into pancreatic endocrine cells and intestine; while cells located two cells away from the midline are inclined to develop into liver and exocrine pancreas (53, 54). The medio-lateral patterning of cell fate decision relies on mesoderm-derived Bmp2b (53). Higher level of Bmp2b promotes liver versus pancreas development while notochord-derived nog2, a Bmp antagonist, increases the number of pancreatic progenitors (55). Given that *id2a* belongs to the inhibitor of differentiation protein family and is a downstream gene of Bmp signaling, we use both *id2a* iCre and CreERT2 knock-in lines for lineage tracing, in order to have a better understanding of the hepatic-pancreatic development. The results from *TgKI(id2a-p2A-mNeonGreen-t2A-iCre);Tg(ubi:CSHm)* demonstrated an universal labelling in the liver and pancreas, suggesting that *id2a^+^* cells are common hepatic-pancreatic multipotent progenitors (Figure 5H and I). Furthermore, we did temporal labelling by treating with 4-OHT at several timepoints (20, 24, 32 and 48 hpf) for 24 hours (Figure 6A-D). Both hepatocytes and hepatic ducts were H2BmCherry positive when labelled at 20 hpf (Figure 6A). However, the H2BmCherry positive cells in the liver were gradually restricted to the hepatic duct at 24 hpf (Figure 6B) and demonstrated specific hepatic duct labelling when the treatment started at 32 hpf or 48 hpf (Figure 6C and D, Figure EV13). In the zebrafish pancreas, similar to *nkx6.1*, temporal labelling of *id2a^+^* cells at 1 dpf marked mainly ductal cells (intra-and extra-pancreatic ductal cells) and a subpopulation of endocrine cells (Figure 6E and F). These results suggest that Bmp signaling is active in liver, dorsal bud-derived pancreas, and ventral bud-derived pancreas specification at early developmental stages; while its activity is mostly maintained in hepatic and pancreatic duct.

**Figure 6.**
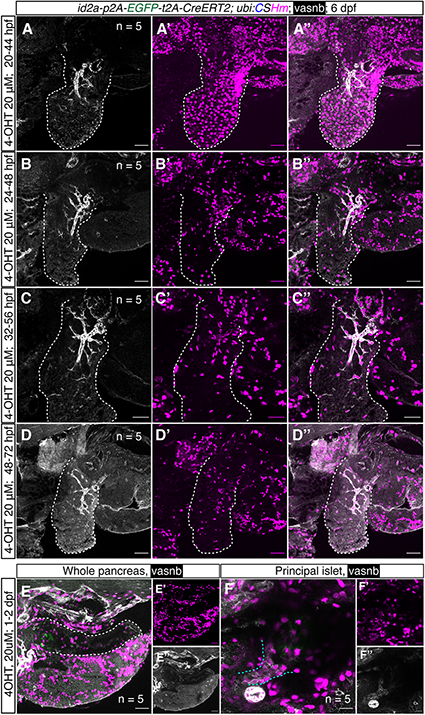
Temporally controlled *id2a* lineage tracing experiment in the zebrafish liver and pancreas. (A-D) Representative confocal images of the liver in *TgKI(id2a-p2A-EGFP-t2A-CreERT2)* treated with 20 μM 4-OHT at 20 hpf (A-A’’), 24 hpf (B-B’’), 32 hpf (C-C’’), and 38 hpf (D-D’’) for 24 hours. For each condition, we scanned 5 embryos; for each embryo, 20 to 32 single planes were imaged. The progenies of *id2a^+^* cells were labelled with H2BmCherry. The white dashed lines indicate liver. (E-F) Representative confocal images of zebrafish pancreas and principal islet at 6 dpf treated with 4-OHT 20 μM at 1-2 dpf. (F’ and F’’) Magnified confocal images of F showing zebrafish principal islets. Extrapancreatic ductal cells and principal islet are defined by anti-vasnb antibody (in white) and anti-glucagon antibody (in green), respectively. The white dashed lines indicate pancreas; while the cyan dashed lines indicate extrapancreatic duct. Scale bars = 35 μm (A-D), 40μm (E) or 20 μm (F).

Lastly, we examined whether there is ductal-to-hepatocyte conversion in two liver injury models, as previous studies showed that *tp1^+^* intrahepatic ductal cells can convert to hepatocytes in an extreme hepatocyte ablation condition (56, 57). Here, we changed responder line to *Tg(−3.5ubb:loxP-EGFP-loxP-mCherry) (*abbreviated as *Tg(ubi:Switch))* for the lineage tracing experiment, as the cytoplasmic mCherry can allow us to distinguish the cell type based on the cell morphology (24148620). The *id2a^+^* hepatic ductal cells were first temporally labelled from 2-3 dpf, with subsequent metronidazole (MTZ) treatment using larvae carrying the *fabp10:CFP-NTR* transgene or chemical-induced severe liver injury (acetaminophen 10 mM + 0.5% ethanol for 48 hours, resulting in near 90% hepatocyte ablation) from 3-5 dpf followed by 2 days of regeneration (34). Based on the morphology of the mCherry positive cells and the vasnb co-staining, we confirmed that a large amount of hepatocytes were derived from an *id2a^+^* duct origin after the MTZ/NTR induced liver injury (Figure 7). However, all lineage-traced cells remained in a ductal identity after the chemical-induced injury (Figure EV14), indicating that *id2a^+^* duct-to-hepatocyte conversion mainly occurs after the extreme hepatocyte loss while this phenomenon is very sparse after the severe liver injury model. Thus, the Notch-and BMP-responsive hepatic ductal cells maintain progenitor potential in zebrafish.

**Figure 7.**
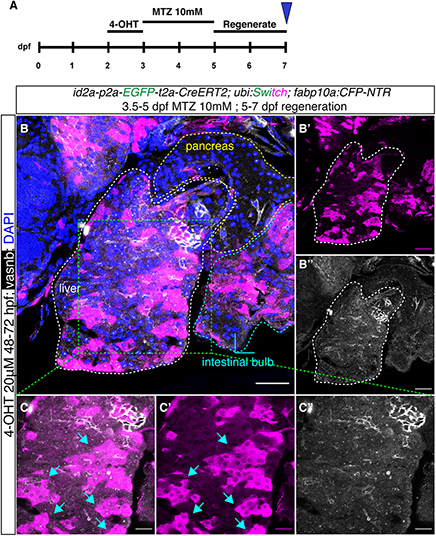
*id2a* lineage-traced cells in an extreme liver injury model. (A) Experimental timeline of *id2a* lineage tracing in a MTZ/NTR induced extreme liver injury model. (B) Representative confocal images of *id2a* lineage-traced cells in the extreme liver injury model. In total, we scanned 5 samples with 21-35 single planes in each zebrafish larvae. The white dashed lines indicate the liver; the yellow dashed lines indicate the pancreas, while the cyan dashed lines indicate the intestinal bulb. (C) The magnified image of the liver showing large numbers of regenerated hepatocytes lineage traced back to an *id2a^+^* cellular origin. The arrows point to clusters of regenerated hepatocytes. Scale bars = 40 μm (B) and 20 μm (C).

## Discussion

Here, we introduce a straightforward 3’ knock-in pipeline to generate zebrafish lines for both cellular labelling and lineage tracing. This method combines a one-step PCR amplification for 5’ modified dsDNA donor with short or long HAs and co-injection with Cas9 protein/gRNA RNPs. Notably, we observed a very high germline transmission rate, indicating that this method can achieve early genetic integration. By systematic comparisons with different donors, we proposed that 5’ modified dsDNAs with short HAs had the best performance when gRNAs spanning over stop codon are available. Lastly, we managed to knock-in large DNA fragments with multiple cassettes linked by 2A peptides and the generated zebrafish lines that can be used for multiple applications. In all, this straightforward and highly efficient knock-in pipeline is versatile and amenable to a wide range of users, allowing researchers to carry out knock-in projects in a scalable fashion.

The CRISPR-Cas9 mediated knock-in method for experimental animals was first introduced in mouse models, and afterwards, optimizations have been developed in various organisms. However, each optimization approach needs to be rigorously tested in each individual model due to their huge differences in early embryonic development. The NHEJ-guided 5’ knock-in upstream of ATG, NHEJ-guided intron-based knock-in, exon-based knock-in (such as the Geneweld toolbox), and 3’ knock-in with circular plasmids have been reported to generate several reporter lines and floxed lines. Nevertheless, the preparation of the donor plasmid is time-consuming and requires multiple steps of molecular cloning. Additionally, the relatively large size and slow diffusion into the nucleus, e.g. with donors in the form of circular plasmids or plasmids with *in vivo* linearization sites, could delay the integration and reduce the mosaicism. This needs to be particularly considered in zebrafish, as the blastomeres divide extremely fast during the cleavage period and such a delay can lead to declined germline transmission (58, 59). Several recent studies used dsDNA as the direct donor. The Tild-CRISPR method, which is based on *in vitro* vector linearization by PCR amplification or enzyme-cutting, can dramatically increase the knock-in efficiency in mice (5). In zebrafish, however, there are conflicting results: one former study showed that such *in vitro* linearization was inefficient to mediate HDR compared with circular plasmid; whereas another study reported that co-injection of synthetic gRNA and linear dsDNA together showed promising results (6, 60). Additionally, PCR amplicons containing fluorescence tagging with several hundred base pairs flanking sequences can be successfully knocked-in into non-coding regions at N or C terminal regions, indicating that PCR amplicons can be useful donors in facilitating zebrafish HDR-based knock-in (10). Interestingly, one study systematically compared 13 modifications on dsDNA donors with short HAs in generating knocked-in human cell lines and showed several modifications (especially 5’ C6-PEG10, amine group with a C6 linker [5’ AmC6] or C12 linker [5’ AmC12]) can exceptionally enhance the 3’ knock-in rate (with up to 500%) (27). Although the underlying mechanism is not fully understood, it is proposed that the 5’ end protection can prevent the NHEJ event and donor multimerization (61).

Here, we focused on 3’ knock-in because this method, theoretically, can keep the endogenous gene functional and intact. We observed that short HAs had extraordinary performance and were even better than long HAs when knocking in long DNA fragments, i.e. when there is a good gRNA spanning over the stop codon region. Furthermore, our data targeting the *krt92* and *krt4* loci are the first evidence suggesting that the end protection greatly improves the efficiency of HDR integration in zebrafish, consistent with that end protection of dsDNA limits exonuclease degradation and concatenation. Given that the dsDNAs with short HAs are smaller in size (compared with circular plasmid or dsDNA with long HAs), we assume they can locate into nucleus in a more efficient way. Altogether, the 5’ end protection and short HAs may lead to a high concentration of dsDNA in the local integration region for the promotion of HDR. It is difficult to perform head-to-head comparisons regarding efficiency between various knock-in methods due to that different loci were chosen for testing (27003937, 25411213, 32064511, 33462297). However, our method achieved an overall higher percentage of F0 mosaicism rate with a stable high percentage of germline transmission rate (equal or more than 50%).

The main advantages of our method become apparent in several aspects. First, we are targeting the 3’ end without the disruption of the endogenous gene products. In addition, the 3’ in-frame integration did not show any effects on the mRNA expression of the endogenous genes (Figure EV15). This is of great importance in developmental and regenerative studies as in certain cases the loss of one gene allele (i.e. in transcription factors) can generate detectable phenotypes (63). Second, we found that dsDNAs with 5’ modifications were efficient donors in zebrafish. As the short HAs were at least as equally good as long HAs in the locus we tested, the preparation of donors can be dramatically simplified by one-step PCR using primers harboring 29-41 base pairs overhangs, which is in line with previous work (14). Third, the design of 2A peptides linking different functional cassettes enables us to generate knock-in lines for multiple uses. Fourth, it is much easier to identify the mosaic founders for knock-in of non-fluorescent protein (e.g. Cre) when combined with a fluorescent protein as an in-frame marker. Fifth, although previous studies supported the use of Cas9 mRNA in medaka (61, 62), here, we instead recommend injecting Cas9 protein in zebrafish embryos to enable early integration. Medaka has a much slower cell cycle during early development (16-cell stage at 3 hpf) (64) while zebrafish embryos display very rapid cell division (over 1000 cells at 3 hpf) (65). The generation of double-stranded DNA break needs to be concordant with a sufficient amount of donor templates ready for HDR; otherwise, cells are inclined to use error-prone NHEJ, which would introduce indels. As Cas9 protein can be transported into the nucleus rapidly, it can cleave the genomic DNA soon after injection, which is particularly important for obtaining high germline transmission in zebrafish. Given that dsDNAs are in smaller sizes and can diffuse faster than whole plasmids, the cleaved genomic DNA has a higher chance to be precisely repaired by MMEJ rather than NHEJ. Sixth, we chose the strongest green fluorescence monomer, mNeonGreen, as an alternative to EGFP for cellular labelling. Strong fluorophores are helpful for the identification of mosaic embryos in case the targeted gene has a low expression. Moreover, the mNeonGreen is not recognized by the anti-GFP antibody, allowing the users to combine it with transgenic lines expressing GFP or its derivatives.

We shall also note that the use of short HAs is highly dependent on whether there is a good gRNA spanning over the stop codon region. If there is an absence of such gRNA, long HAs might be a better choice as mutations need to be introduced in the left (for gRNA site upstream of stop codon) or right (for gRNA site downstream of stop codon) HAs, such that the gRNA does not also cleave the donor dsDNA. However, the use of short HAs provides an easy and efficient way to knock-in a specific donor into several candidate genes in parallel, i.e. without needing to perform complex molecular cloning steps.

We also employed our knock-in lines to investigate the cell fate determination in the liver and pancreas. We, for the first time, used a lineage-tracing method to make a definitive conclusion that the *nkx6.1^+^* intrapancreatic ductal cells can serve as progenitor cells that can differentiate into endocrine cells in secondary islets. Also, by combining the lineage tracing results from non-inducible and inducible Cre lines, we depicted a temporal cell differentiation map for the specification between the acinar fate and ductal/endocrine fate in the pancreas, as well as between hepatocyte fate and hepatic ductal fate in the liver. Lastly, we consolidated and extended the previous findings indicating that the Notch-and BMP-responsive liver ductal cells are progenitors and able to convert to hepatocytes only upon extreme liver injury (24148620, 24315993). Further experiments using lineage tracing, single-cell RNA sequencing and tissue specific gain-and-loss of function are warranted to investigate detailed cellular and molecular mechanisms in the development and regeneration of these tissues.

Last but not least, by systematic comparisons of the *krt4* gene expression pattern in both widely used transgenics and our knock-in lines, we found that our knock-ins can fully recapitulate the endogenous gene expression; while the *krt4 cis*-regulatory element cloned for skin cell labelling is incapable to drive the reporter gene expression in the intestine (intestinal bulb and hindgut). This is of particular importance for the lineage-related research as most previous cell lineage discoveries in zebrafish are based upon cloning promoters for transgenics, which may fail to label certain cell types, display ectopic expression or exhibit leakage issues. This might be due to the fact that different tissues/cell types tend to use different *cis*-regulatory elements and chromatin structure, as well as the enhancer-promoter loops might have remarkable differences in various cell types (25650801). It is really difficult to precisely predict the exact region of the regulatory sequences sufficient to activate the gene expression in each cell type without a comprehensive profiling of the epigenetic landscape in a single-cell resolution. Therefore, we believe that our method, together with other zebrafish knock-in Cre/CreERT2 methods, might divert the standard toward the knock-in based genetic fate mapping for both confirming old and making new discoveries.

In summary, we described a novel 3’ knock-in pipeline for the construction of zebrafish lines with multiple cassettes. This method is easy to implement as it only includes a one-step PCR reaction and co-injection with Cas9/gRNA RNPs. The application of dsDNA with short HAs allows us to knock-in specific donors in multiple genes in a scalable fashion. This method is highly efficient as it can achieve a desirable percentage of germline transmission from mosaic F0, which is helpful for small-to-medium sized zebrafish labs with limited space to screen for founders. The design using 2A peptides for linkage makes it possible to knock-in multiple cassettes at the same genetic locus, further expanding the utility of the knock-in lines. Therefore, we anticipate this efficient and straightforward knock-in method will be of widespread use in the zebrafish field.

## Materials and Methods

### Zebrafish lines used in the study

Males and females ranging from 3 months to 2 years were used for breeding. Zebrafish larvae were incubated in 28.5 °C until 7 dpf. The following published transgenic zebrafish (*Danio rerio*) lines were used in this study: *Tg(ptf1a:GFP)^jh1^* (28), *Tg(Tp1bglob:H2BmCherry)^S939^* (29) abbreviated *Tg(Tp1:H2BmCherry)*, *Tg(−3.5ubb:loxP-EGFP-loxP-mCherry)^cz1701^* (30) abbreviated as *Tg(ubi:Switch)*, *Tg(UBB:loxP-CFP-STOP-Terminator-loxP-hmgb1-mCherry)^jh63^* (31) abbreviated as *Tg(ubi:CSHm)*, *Tg(ela3l:H2BGFP)* (32), *Tg(krt4:EGFP-Mmu.Rpl10a)* (24367659) and *Tg(fabp10a:CFP-NTR)^S931^*.

The following lines were newly generated by the CRISPR-Cas9 3’ knock-in strategy: *TgKI(krt92-p2A-EGFP-t2A-CreERT2)^KI126^*, *TgKI(krt4-p2A-mNeonGreen)^KI127^*, *TgKI(krt4-p2A-mNeonGreen-t2A-iCre)^KI128^*, *TgKI(krt4-p2A-EGFP-t2A-CreERT2)^KI129^*, *TgKI(nkx6.1-p2A-mNeonGreen)^KI130^*, *TgKI(nkx6.1-p2A-mNeonGreen-t2A-iCre)^KI131^*, *TgKI(nkx6.1-p2A-EGFP-t2A-CreERT2)^KI132^*, *TgKI(id2a-p2A-mNeonGreen)^KI133^*, *TgKI(id2a-p2A-mNeonGreen-t2A-iCre)^KI134^* and *TgKI(id2a-p2A-EGFP-t2A-CreERT2)^KI135^*.

Adult fish were maintained on a 14:10 light/dark cycle at 28°C. All studies involving zebrafish were performed in accordance with local guidelines and regulations, and approved by regional authorities.

### The vector design for 3’ knock-in

The vector templates were manually designed for *krt92* and *nkx6.1* 3’ end loci. The vectors include a left long HA of 950 base pairs genomic sequence upstream of the stop codon of the endogenous gene product followed by a GSG linker (glycine-serine-glycine, GGAAGCGGA), p2A sequence (GCTACTAACTTCAGCCTGCTGAAGCAGGCTGGAGACGTGGAGGAGAACCCTGGACCT), zebrafish codon optimized EGFP or mNeonGreen (without stop codons), GSG linker, t2A sequence (GAGGGCAGAGGCAGTCTGCTGACATGCGGTGATGTGGAAGAGAATCCCGGCCCT), zebrafish codon optimized iCre or CreERT2 (with stop codons) and 950 base pairs right long HA downstream of endogenous stop codon flanked by *in vivo* linearization sites (GAGCTCGGTACCCGGGGATC[AGG] on the left; ATCCTCTAGAGTCGACCTGC[AGG] on the right).

### PCR amplification and gel purification

The 5’ modified primers were ordered with AmC6 5’ modification from Integrated DNA Technologies (IDT). The primer powders were diluted with distilled water into 100 mM as stock solution. The 50 μL PCR mixture include:

Forward primer: 2.5 μL
Reverse primer: 2.5 μL
Template plasmid: 1 μL
Distilled water: 19 μL
Q5 Hot Star high-fidelity 2× master mix: 25 μL

We use the following PCR cycle setting:

Pre-denaturing: 98 °C, 30 sec
Denaturing: 98 °C, 10 sec
Annealing: 58-60 °C, 20 sec
Extension: 72 °C, 90 sec
Final extension: 72 °C, 2 minutes and hold on 4 °C

Next, we ran the PCR products in 1% agarose gel with 100V for 45-60 minutes. The corresponding bands were cut out and purified using wizard SV gel and PCR clean-up system (Promega, A9282). The concentrations of purified PCR products were measured by Nanodrop 2000c and then diluted into 70-100ng/uL with distilled water accordingly. The purified PCR products were stored in −20 °C before injection.

### The selection of gRNA

We used the CHOPCHOP web-based tool (http://chopchop.cbu.uib.no/) and set the reference genome as “danRer11/GRCz11”. We selected “CRISPR/Cas9” and “knock-in” module after determining the targeted gene. This tool would scan through the gene exon regions and rank the gRNA based on efficiency score, self-complementarity, and the number of mismatches. Apart from the *in silico* prediction, we also manually examined the 3’ end in the Ensembl database (https://www.ensembl.org/Danio_rerio/Info/Index) to avoid polymorphisms in the targeting site. The following gRNAs were used in this study (with PAM sequence in the brackets):

*krt92* (on reverse strand): 5’-AACCTCGCTCGAGATTGGG(AGG)-3’
*krt4* (on forward strand): 5’-GTCAGCAGTAAACGCTATT(AGG)-3’
*nkx6.1* (on forward strand): 5’-AGAGCTCGTAAAAAGGAAAC(GGG)-3’
*id2a* (on forward strand): 5’-AGGACACTTTACCGTTAATC(AGG)-3’

For the control experiment using plasmid with linearization sites, we used the following gRNAs:

Left: 5’-GAGCTCGGTACCCGGGGATC(AGG)-3’
Right: 5’-ATCCTCTAGAGTCGACCTGC(AGG)-3’

### In vitro pre-assembly of gRNA, Cas9 protein and donor dsDNA

The chemically synthesized Alt-R-modified crRNA, tracrRNA, high-fidelity (Hifi) Cas9 protein, and nuclease free duplex buffer were ordered from IDT. The crRNA and tracrRNA powder were diluted into 100 μM with nuclease free duplex buffer. The 10 μL Hifi Cas9 protein were aliquoted into 10 tubes followed by 1:5 dilution with Opti-MEM (ThermoFisher Scientific, 31985062) solution before use.

Next, we prepared 10 μM crRNA:tracrRNA duplex solution by mixing 1μL crRNA stock solution, 1μL tracrRNA stock solution and 8 μL nuclease free duplex buffer and then incubated in PCR machine, in 95 °C for 3 minutes followed by natural cooling in room temperature for 15 minutes. Afterwards, we prepared the Cas9/gRNA RNP by mixing 2 μL Cas9 protein solution and 2 μL crRNA:tracrRNA duplex solution in 37 °C for 10 minutes. Lastly, we mixed 2 μL Cas9/gRNA RNP, 5 μL donor dsDNA and 0.8 μL phenol red (sigma, P0290) and stored in 4 °C. We recommend to perform this pre-assembly step the day before injection.

### Microinjection and sorting for mosaic F0

We injected 1-2 nL Cas9 RNP and donor dsDNA (50-70 pg/nL) into zebrafish embryos at early one-cell stage. The overall mortality rate was around 50% and we sorted out all dead embryos in the following days. We selected mosaic F0 at 2 dpf based on the fluorescence in the skin (*krt92* and *krt4*), hindbrain and spinal cord (*nkx6.1*) and hindbrain, spinal cord, and olfactory organ (*id2a*) under a widefield fluorescence microscope LEICA M165 FC (Leica Microsystems) using either the GFP (EGFP or mNeonGreen) or YFP (mNeonGreen) channel. Positive mosaic F0 were put into the fish facility at 6 dpf.

### Genotyping of F1

The clipped zebrafish fins were added to lysis buffer (10mM Tris-HCl pH 7.5, 1mM EDTA, 50MM KCL, 0.3% TWEEN 20) and boiled at 95 °C for 10 min. Next, we added 10% volume proteinase K (10mg/ml) and incubated in 55 °C overnight. On the following day, we heat-inactivated proteinase K by boiling at 95 °C for 10 min and used 1 uL as the template for PCR reactions. The following primers were used to amplify the fragments over the insertion site, and the reverse primers were used as the sequencing primer:

*krt92* (forward primer): 5’-CAAGCTCAAGCTCAAGTTCC-3’
*krt4* (forward primer): 5’-GTTATGGTGGTAGCGGCTCTGG-3’
*nkx6.1* (forward primer): 5’-CGACGACGACTACAATAAACC-3’
*id2a* (forward primer): 5’-CTCGACTCCAATTCGGCG-3’
EGFP (reverse primer): 5’-CATGTGGTCGGGGTAGCG-3’
mNeonGreen (reverse primer): 5’-ACTGATGGAAGCCATACCCG-3’

### Quantitative RT-PCR

qPCR was performed using SYBR Green on a ViiA 7 Real-time PCR machine. The gene encoding β-actin was used as the control for normalization. The primer sequences are as follows:

*krt92* (forward primer): 5’-CCGAAACCCTCACCAAGGAA-3’
*krt92* (reverse primer): 5’-CCTCGCTCGTAGATTGGGAG-3’
*krt4* (forward primer): 5’-AACAAGCGTGCTTCCGTAGA-3’
*krt4* (reverse primer): 5’-GCGATCATGCGGTTGAGTTC-3’
*nkx6.1* (forward primer): 5’-CGTGCTCACATCAAAAC-3’
*nkx6.1* (reverse primer): 5’-CGGTTTTGAAACCACACCTT-3’
*id2a* (forward primer): 5’-CAGATCGCGCTCGACTCCAA-3’
*id2a* (reverse primer): 5’-CAGGGGTGTTCTGGATGTCCC-3’
*β-actin* (forward primer): 5’-CGAGCAGGAGATGGGAACC-3’
*β-actin* (reverse primer): 5’-CAACGGAAACGCTCATTGC-3’

qPCR data are expressed as the fold change (ΔΔCt). Four zebrafish larvae were pool together as a biological replicate. We have four biological replicates for each knock-in and the wildtype. Two-tailed t-test was used with p-value = 0.05 as statistical significance.

### Lineage tracing by tamoxifen-inducible Cre recombinase

We used both iCre and CreERT2 lines for the testing of Cre and genetic lineage-tracing experiment. The knock-in iCre lines were crossed with either *Tg(ubi:Switch)* or *Tg(ubi:CSHm)*. For temporal labelling, we treated the zebrafish larvae carrying knock-in CreERT2 as well as *ubi:CSHm or ubb:Switch* transgene with 20 μM 4-OHT (Sigma-Aldrich) in E3 medium in 24-well plates, with 4-8 embryos/larvae per well, for 24-36 hours without refreshment. Upon induction by 4-OHT, cytoplasmic CreERT2 would be translocated into the nucleus to excise the DNA in between the two loxp sites to enable downstream mCherry or H2BmCherry expression.

### Sample fixation for immunostaining

Before fixing the zebrafish larvae/juveniles, we confirmed the presence of the transgenes or the sorting markers by examining the corresponding fluorescence signals under LEICA M165 FC fluorescence microscope. We euthanized the zebrafish juveniles with 250 mg/L tricaine (Sigma-Aldrich) in E3 medium. Before fixation, we washed the zebrafish larvae/juveniles with distilled water three times. We fixed the samples in 4% formaldehyde (Sigma-Aldrich) in PBS (ThermoFisher Scientific) at 4 °C for at least 24 hours. Following three washes with PBS, we removed the skin and crystallized yolk (of the zebrafish larvae) by forceps under the microscope to expose the pancreas and liver for immunostaining.

### Confocal imaging of live zebrafish larvae

We prepared 1% low-melting agarose gel by dissolving 0.01 g agarose (sigma, A9414) in 1ml E3 solution with 250 mg/L tricaine followed by 65 °C heating for 10 min and cooling on a wedged zebrafish mold. Zebrafish larvae were euthanized in E3 medium with 250 mg/L tricaine and then repositioned in the gel groove to a suitable position. The confocal imaging was performed using the laser scanning microscopy platform Leica TCS SP8 (Leica Microsystems) with a 10× objective.

### Immunostaining and confocal imaging

We performed immunostaining similarly to our previous report (33). In brief, we firstly performed blocking step by incubating the zebrafish samples in blocking solution (0.3% Triton X-100, 4% BSA in PBS) at room temperature for at least one hour. We then incubated the samples in blocking solution with primary antibodies at 4 °C overnight. After removing the primary antibodies, we washed the samples with washing buffer (0.3% Triton X-100 in PBS) ten times at room temperature for at least four hours. Next, we incubated the samples in blocking solution with fluorescent dye-conjugated secondary antibodies and the nuclear counterstain DAPI (ThermoFisher Scientific) at 4 °C overnight. Afterwards, we removed the secondary antibodies and DAPI and washed the samples with washing buffer ten times at room temperature for at least four hours. The following primary antibodies were used: chicken anti-GFP (1:500, Aves Labs, GFP-1020), goat anti-tdTomato (1:500, MyBioSource, MBS448092), mouse anti-mNeonGreen (1:50, Chromotek, 32F6), rabbit anti-insulin (1:100, Cambridge Research Biochemicals, customised), mouse anti-glucagon (1:50, Sigma, G2654), rabbit anti-cdh17 (1:1000, customised sera, gift from Prof. Ying Cao, Tongji University) and rabbit anti-vasnb (1:1000, customised sera, gift from Dr. Paolo Panza).

Before confocal imaging, we mounted the stained samples in VECTASHIELD Antifade Mounting Medium (Vector Laboratories) on microscope slides with the pancreas or liver facing the cover slips. We imaged the pancreas and liver with the Leica TCS SP8 platform.

### Hepatocyte ablation by chemo-genetic and pharmacological approaches

The extreme hepatocyte injury model was induced based on metronidazole/nitroreductase (MTZ/NTR) system (18536643). In brief, the *TgKI(id2a-EGFP-t2a-CreERT2); Tg(ubb:Switch); Tg(fabp10a:CFP-NTR)* larva were treated with 10 mM MTZ (final concentration)_in E3 medium (24148620). The severe hepatocyte injury model included incubating the zebrafish larvae in E3 medium supplemented with 10 mM acetaminophen (Sigma-Aldrich, A7085), and 0.5% Ethanol for 48 hours from 3 to 5 dpf, where after the larvae were washed three washes with E3 solution and allowed to recover for 2 days, as previously reported (34).

### Statistical analysis and data visualization

Similar experiments were performed at least two times independently. The number of cells in the confocal microscopy images were all quantified manually with the aid of the Multipoint Tool from ImageJ. The scheme of knock-in strategy was illustrated by using “IBS” software (35). Statistical analyses were carried out by two-tailed *Mann Whitney U* tests (comparing two groups) or by *Kruskal-Wallis* tests (comparing three groups) unless otherwise stated. The results were presented as the mean values ± SEM and *P* values ≤ 0.05 are considered as statistically significant. The *n* number represents the number of zebrafish in each group for each experiment. All statistical analyses and data visualization were performed on R platform (version 4.0.2) using “ggplot2” and “ggpubr” packages.

### Data availability

The key resource table, the construct maps and the Sanger sequencing files that support the findings of this study are available in the public depository (https://osf.io/tdkvh/).

## Acknowledgements

We thank Dr. Ka-Cheuk Liu for critical reading of the manuscript. We also thank Chikai Zhou (Karolinska Institutet), Kazuyuki Hoshijima (University of Utah), Remy Manuel (Uppsala University), Qingshun Zhao (Nanjing University), Maura McGrail (Iowa State University), and Pierre Gillotay (ULB) for the informative discussion, Ying Cao (Tongji University) and Paolo Panza (Max Planck Institute for Heart and Lung Research) for their antibodies as gifts and Donghun Shin (University of Pittsburgh), Nikolay Ninov (CRTD Dresden) and Elke Ober (University of Copenhagen) for sharing the fishlines. The illustration of the double-stranded DNA and tracr:crRNA complex in Figure 1 was created by Biorender.com. Research in the lab of O.A. was supported by funding from the European Research Council under the Horizon 2020 research and innovation programme (grant n° 772365); the Swedish Research Council; the Novo Nordisk Foundation; and the Strategic Research Programmes in Diabetes at the Karolinska Institutet. J.M. was supported by China Scholarship Council (CSC).

